# Non-negative least squares computation for in vivo myelin mapping using simulated multi-echo spin-echo T_2_ decay data

**DOI:** 10.1101/2020.02.15.948976

**Authors:** Vanessa Wiggermann, Irene Vavasour, Shannon Kolind, Alex MacKay, Gunther Helms, Alexander Rauscher

**Affiliations:** Department of Physics and Astronomy, University of British Columbia; Department of Pediatrics, University of British Columbia; UBC MRI Research Centre, University of British Columbia; Department of Radiology, University of British Columbia; Department of Medicine, University of British Columbia; Department of Clinical Sciences Lund, Lund University

**Keywords:** myelin, T_2_ distribution, T_2_ decay, relaxation, non-negative lease squares, flip angles, noise, spin echo

## Abstract

Multi-compartment T_2_-mapping has gained particular relevance for the study of myelin water in brain. As a facilitator of rapid saltatory axonal signal transmission, myelin is a cornerstone indicator of white matter development and function. Regularized non-negative least squares fitting of multi-echo T_2_ data has been widely employed for the computation of the myelin water fraction (MWF) and the obtained MWF maps have been histopathologically validated. MWF measurements depend upon the quality of the data acquisition, B1^+^-homogeneity and a range of fitting parameters. In this special issue article, we discuss the relevance of these factors for the accurate computation of multi-compartment T_2_ and MWF maps. We generated multi-echo spin-echo T_2_ decay curves following the approach of CarrPurcell-Meiboom-Gill for various myelin concentrations and myelin T_2_ scenarios by simulating the evolution of the magnetization vector between echoes based on the Bloch equations. We demonstrated that noise and imperfect refocusing flip angles yield systematic underestimations in MWF and intra-/extracellular water geometric mean (gm) T_2_. MWF estimates were more stable than myelin water gmT_2_ time across different settings of the T_2_ analysis. We observed that the lower limit of the T_2_ distribution grid should be slightly shorter than TE1. Both TE1 and the acquisition echo spacing also have to be sufficiently short to capture the rapidly decaying myelin water T_2_ signal. Among all parameters of interest, the estimated MWF and intra-/extracellular water gmT_2_ differed by approximately 0.13-4 percentage points and 3-4 ms, respectively, from the true values, with larger deviations observed in the presence of greater B1^+^-inhomogeneities and at lower signal-to-noise ratio. Tailoring acquisition strategies may allow to better characterize the T_2_ distribution, including the myelin water, in vivo.

---

Since the inception of magnetic resonance imaging (MRI), researchers have strived to use MR signal relaxation techniques for the characterization of tissues and tissue pathologies. T_2_-weighting by means of spin-echoes, first described by Erwin Hahn [1,2], captures the effect of random field fluctuations on the transverse MR signal. Quantitative T_2_ relaxation measurements are commonly performed using the Carr-Purcell-Meiboom-Gill (CPMG) acquisition scheme [3,4]. Rapid acquisitions or 3D encoding utilize a “train” of spin-echoes, a technique pioneered by Jürgen Hennig [5]. Hennig also devised the extended phase graph (EPG), used to describe the formation of spurious echoes in the T_2_ decay due to imperfect refocusing flip angles (FA) in the presence of static field inhomogeneities [6,7]. By ensuring that subsequent refocusing pulses have the same phase, orthogonal to that of the excitation pulse, CPMG mitigates errors due to imperfect refocusing and achieves higher signal through constructive interference of refocusing pathways. T_2_-mapping has been primarily applied to characterize brain tissue [8,9], but also outside the brain for imaging of cartilage [10] and intervertebral discs [11]. Notably, microstructural tissue compartmentalization can contribute more than one exponential component to the T_2_ decay [12–15].

The short T_2_ component in the brain and its relationship with myelin Multi-component analysis of multi-echo T_2_ data through construction of continuous T_2_ distributions was described by Whittall and MacKay [16]. They employed non-negative least squares (NNLS) [17] to solve the inverse problem that is posed by an unknown number of T_2_ components contributing to the T_2_ decay. In the early nineties, Menon and Allen [18] described two T_2_ components in feline brain ex vivo and assigned the shorter component to water in myelin. Thereafter, a similar T_2_ component was observed in guinea pig brain ex vivo [13]. MacKay and Whittall et al. were the first to report a short T_2_ component in human brain in vivo at 1.5 T [19]. This short T_2_ signal co-localized with brain white matter (WM) and its signal proportion was in line with the amounts of “myelin water” (MW) reported in vitro [20]. These observations strongly suggested that the short T_2_ signal originates from water molecules trapped within the tightly wrapped myelin bilayers. At 30 Å [21], i.e. only about ten times the size of water molecules, the space occupied by water within the bilayers is much narrower than intra- and inter-axonal distances. The therefore restricted motion leads to faster spin-spin dephasing of aqueous protons in MW than in the intra- and extracellular space. The MR signal from non-aqueous protons of the phospholipid bilayers decays to zero in less than 100 *μ*s; hence it is not visible with conventional MR systems [22]. The relationship of the short T_2_ component and myelin was illustrated in cases of known myelin pathology, where lesions devoid of myelin showed little short T_2_ signal [23,24]. Histopathological studies later validated the association of the short T_2_ component with myelin lipid concentrations [25,26]. The term myelin water fraction (MWF) was coined for the short T_2_ fraction of the total water signal. However, water protons other than those in myelin may contribute to rapid T_2_ decay. Biological iron, calcium and other tissue constituents can cause additional T_2_ shortening of water signal in the intra- and extracellular space [27], increasing the apparent MWF [28]. Since WM iron is predominantly found in myelin forming oligodendrocytes and contributes to lipid synthesis [29], the sensitivity of the short T_2_ component to both myelin and iron should be considered. Mapping myelin in vivo permits us to study brain development [30–32], trauma [33] or aging [34,35], as well as neurodegenerative diseases that have a direct link to myelination, such as multiple sclerosis (MS) [23,36]. However, despite being proposed 25 years ago, spin-echo myelin water imaging (MWI) has to-date found limited clinical application. This is in part due to the long acquisition times and an intricate signal analysis.

MWI using non-negative least squares (NNLS) fitting to multi-exponential T_2_ data The NNLS analysis and the computation of the MWF depend on a number of parameters. In vivo literature has primarily reported on multi-component T_2_-mapping and corresponding analysis parameters in human brain at 1.5 T and 3 T. As MWI expands to other field strengths and applications for which multi-component T_2_ times have not yet been established, adjustment of the T_2_ analysis and application of NNLS may be required to obtain reliable MWF measurements. For example, if T_2_ relaxation times are shorter, the T_2_ boundaries that distinguish the different water compartments need to be redefined. Otherwise, signal from a water pool with a T_2_ longer than that of MW may be erroneously counted toward the short water pool fraction as it crosses the pre-established T_2_ cut-off [37]. In brain tissues, the intermediate water compartment is attributed to intra- and extracellular water (IEW). This intermediate T_2_ pool can typically not be further divided using the resolution of T_2_ time at 3 T. However, a few studies have found two T_2_ times in the intermediate T_2_ range, ascribing the longer component to extracellular water [38–40].

To provide greater insight into the data analysis, we discuss in this article the various parameters involved in the computation of the NNLS solution [17] in accurately determining different water pool fractions and their T_2_ properties. A number of groups have assessed the properties of NNLS for multi-component T_2_ analysis, particularly in the first decade of MWI [16,41,42–44]. We intend to update those findings and provide a guideline for the application of NNLS with EPG to multi-component T_2_ analysis in different settings. The main challenge for testing analysis parameters in vivo relates to the lack of a known ground truth. The MWF and even more so the T_2_ of myelin water (MW) are not well established. Differences in acquisition timing may further influence the range of T_2_ time that is accessible for analysis of T_2_ decay data [41]. In this work, we used Bloch simulations to generate signal decay curves from different multi-echo T_2_ acquisition strategies for synthetic voxels that contained realistic myelin geometries. First, the EPG algorithm was employed to estimate the unknown refocusing FAs [7,45]. Thereafter, the generated T_2_ decay curves were analyzed with varying NNLS parameters and results were compared to the known ground truth. We addressed the influence of noise and regularization on the computation, in scenarios of low to high MWFs and different MW T_2_ times.

## Experimental

### Numerical simulation of T2 decay data

Bloch simulations were performed on a 256 × 256 grid over a 2D voxel (Figure 1). The layout was based on a publicly available electron microscopy image (https://upload.wikimedia.org/wikipedia/commons/c/c1/Myelinated_neuron.jpg, accessed Sept 12th 2018). The voxel contained three myelinated axons, two of which had thinner myelin sheaths than the third, amounting to 21% MWF (Fig. 1A). To assess whether lower MW contents can also be correctly estimated, the voxel layout was adjusted to yield approximately 11.6% and 5.1% MWF (Fig. 1B and C). These models are henceforth denoted as models (M) 1, 2, 3, corresponding to 21, 11.6 and 5.1% MWF.

**Fig. 1.**
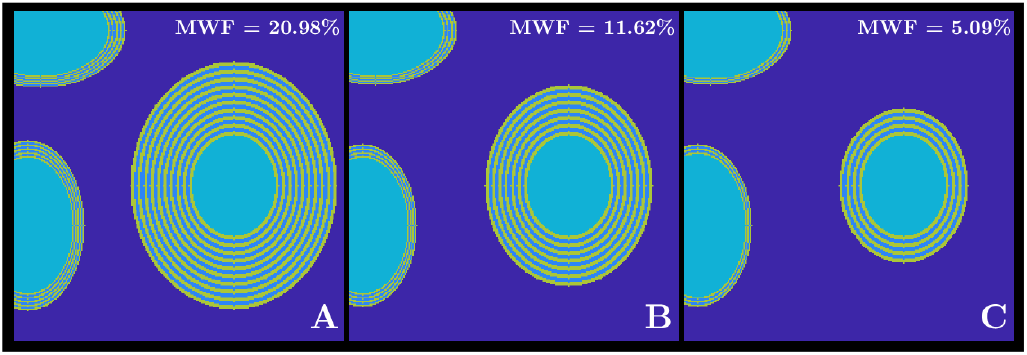
Geometric voxel design utilized for multi-echo spin-echo Bloch simulations. Spins were equally distributed on a 256 × 256 grid over the 2D voxel. The MWF was varied between the three scenarios, yielding approximately 21%, 11.6% and 5.1% for A, B and C, respectively. These are subsequently referred to as Model 1, 2 and 3 (M1, M2, M3), reflecting high to low myelin content. The colouring only distinguishes the four different tissue compartments and does not reflect different assigned relaxation times. Using this layout, tissue relaxation times were assigned to all spins by sampling from a Gaussian distribution around the mean compartmental relaxation time.

We defined four subspaces: Extra-axonal water, intra-axonal water, water in between the myelin bilayers and the non-aqueous myelin bilayers. Literature T_2_ and T_1_ relaxation times [46–48] were assigned to spins in different compartments by random sampling from a Gaussian distribution around the mean compartmental T_1_ and T_2_. Local spin resonance frequencies were allocated by forward field computation of the magnetic susceptibility distribution [49] (*χ* = 0 ppm for all water, *χ*= −0.02 ppm for the myelin lipid bilayers, assuming 80% myelin lipids and 20% proteins [50,51]). Resonance frequency offsets (f) [52–54], relative to the extra-axonal water frequency, were then assigned to the different water pools considering the computed field perturbations. Intra- and extracellular water were assumed to have the same T_1_ and T_2_ (T_1_ = 1200 ± 200 ms [47,48], T_2_ = 70 ± 10 ms [46]), but different f (−3.052 and 0.053 Hz, respectively). MW was presumed to exhibit a rapid signal decay in both T_1_ and T_2_ (T_1_ = 600 ± 100 ms [47,48], T_2_= varied from 3 – 20 ± 5 ms [46], f = 7 Hz [53,54]). Very short, (“MR invisible”) relaxation rates and high offset frequencies were modelled for the highly anisotropic non-aqueous myelin bilayers (T_1_ = 120 ± 20 ms, T_2_ = 0.1 ± 0.05 ms, f =−20 Hz). Note that the proton signal from myelin phospholipid bilayers does not decay exponentially, but is characterized by a Super-Lorentzian line shape [55,56]. Literature on T_1_ of the aqueous and non-aqueous parts of myelin is scarce and in vivo measurements are known to be affected by magnetization exchange [57]. Hence, T_1_ values represent approximate values only.

A linear frequency variation was added to the frequency map as encoding gradients impose much larger frequency offsets than induced locally [7]. Gaussian noise was added to the T_1_ and T_2_ maps and independently to the real and imaginary components of the complex field map (signal-to-noise ratio (SNR) = 250).

Multi-echo spin-echo data were generated analogous to the in vivo MR protocol of the gradient-echo and spin-echo acquisition currently performed for MWI at 3 T [58]: 32 echoes, TE1/ΔTE=10/10 ms and TR=1200 ms. The magnetization vector evolution was described with rotation matrices for each isochromat, allowing for spin excitation and dephasing in the rotating frame of reference. The Carr-Purcell-Meiboom-Gill spin-echo experiments consisted of a 90° excitation pulse along *ŷ* and subsequent 180° refocusing pulses along, with the time between the excitation and first refocusing pulse being TE_1_/2. At time of echo formation, the total magnetization of the set of isochromats was computed and the magnitude of the complex signal, i.e. the transverse component of the magnetization vector, was taken. For each of the three models (Fig. 1), we varied the MW T_2_ through 3, 5, 8, 10, 13, 15, 18 and 20 ms. Six refocusing FAs between 130°−180° were considered and the effect of shorter echo spacing was tested by also computing decay curves of 48 echoes at 8 ms TE/ΔTE and 64 echoes at 6 ms TE/ΔTE [59]. Note that the 48 and 64 echo acquisitions included a few later TE times than the 32 echo acquisition. To accommodate the longer readouts, TR was chosen longer than typically used at 3 T for 32 echoes.

### Computation of T_2_ distributions and analysis parameters of interest

The most fundamental step in correctly estimating T_2_ and subsequently the MWF from multi-echo MRI is the determination of the actual refocusing FA to account for the presence of stimulated echoes [45]. To this end, hypothetical T_2_ decay curves are computed by EPG for a number of T_2_ times (T_2,*i*_) and at a few different FAs, typically eight, equally spaced between 100°−180°. EPG identifies three magnetization phase configurations and traces state transitions by relaxation and transition matrices. The states, F_*n*_, 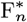 and Z_*n*_, represent transverse and longitudinal dephasing and rephasing spins. Following the application of a radiofrequency pulse *α* with an initial phase of zero, state transitions occur by [45]

**Table 1.** Summary of the RMSE estimates between true and estimated rFAs. RMSE are given in units of [° FA]. M1, M2, M3 indicate the three models of different MWFs, going from highest in M1 (21%) to lowest in M3 (5.1%).

| Sequence \ SNR |  | SNR |  |  |  |  |  |  |  |  |
| --- | --- | --- | --- | --- | --- | --- | --- | --- | --- | --- |
|  |  | 30 | 50 | 100 | 150 | 200 | 250 | 300 | 500 | 1000 |
| M1 | M1 - 32e | 8.74 | 6.82 | 4.02 | 5.77 | 3.24 | 2.38 | 3.40 | 1.39 | 1.48 |
|  | M1 - 48e | 3.81 | 6.03 | 3.30 | 3.39 | 5.18 | 3.98 | 3.79 | 2.61 | 2.42 |
|  | M1 - 64e | 5.80 | 3.07 | 4.87 | 6.59 | 3.73 | 4.71 | 4.74 | 3.91 | 3.35 |
| M2 | M2 - 32e | 8.44 | 8.26 | 4.58 | 6.13 | 3.11 | 2.13 | 2.72 | 3.11 | 2.12 |
|  | M2 - 48e | 7.13 | 5.01 | 4.75 | 5.46 | 4.09 | 2.48 | 2.90 | 3.12 | 2.90 |
|  | M2 - 64e | 4.83 | 5.04 | 5.18 | 5.75 | 4.21 | 4.48 | 2.66 | 4.19 | 3.04 |
| M3 | M3 - 32e | 8.09 | 10.19 | 4.83 | 3.86 | 4.24 | 2.25 | 3.32 | 2.30 | 2.39 |
|  | M3 - 48e | 5.97 | 6.21 | 5.59 | 3.84 | 5.69 | 2.30 | 4.28 | 2.14 | 1.68 |
|  | M3 - 64e | 7.34 | 5.52 | 4.17 | 7.66 | 2.62 | 4.94 | 3.63 | 4.02 | 2.65 |

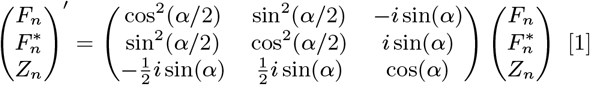

Between refocusing pulses, spins may transition from F_*n*_ to 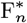 and reversely, noting that the sum of the magnetization in all states has to remain constant. Echoes are formed when 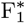 transitions to F_1_, with echo amplitudes corresponding to the magnitude of 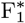. When periodicity conditions are met, i.e. TE_*n*_ = n·ΔTE, the number of configurations that need to be computed grows linearly with the number of echoes. If echoes with different phases occur, additional states need to be traced. An average, single exponential tissue T_1_ (1 s for WM at 3 T [9]) and sequence parameters are also considered by EPG, including echo train length (ETL), Δ TE and the attempted refocusing pulse angle, customarily 180° for spin-echo sequences. Note that computing voxel specific T_1_ times does not improve MWF estimation [45]. For each FA, NNLS determines the corresponding T_2_ distribution. A T_2_ decay curve is then computed by back projection from the T_2_ distribution and compared to the measured data. The sum of squares of the residuals, i.e. *χ*^2^, is calculated for each FA.

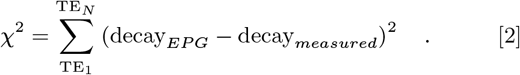

By cubic spline interpolation of all eight *χ*^2^, the FA corresponding to the minimum *χ*^2^, i.e. the actual FA, is established.

With the determined FA, EPG is used to recompute T_2_ decay curves for each T_2*i*_ time in the grid. The magnetization computed at each echo will depend on the echo amplitudes computed by EPG, the assumed proton density (PD), the longitudinal magnetization M0 and the excitation FA [45].

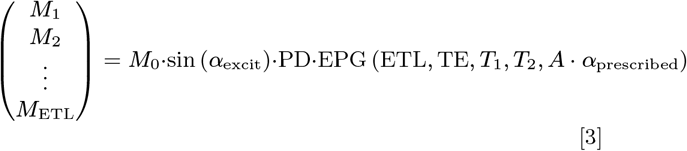

A represents the already computed deviation from the prescribed FA in each voxel. NNLS subsequently estimates the T_2_ distribution by comparing the theoretical decay curves to the measured data. The data remains unaltered when using EPG. Only the generated decay curves are adjusted to reflect imperfect signal refocusing and T_1_ effects in the T_2_ decay, thereby matching better to the acquired data. Due to discretization, resulting T_2_ distributions feature only delta-peaks at few T_2_ times, which may vary for different noise realizations. The sum S(T_2_) of the delta-functions for a given T_2_ with amplitudes ai determined at times T_2*i*_ can be expressed as:

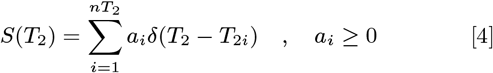

The amplitudes ai are larger or equal zero as T_2*i*_ times are either present with a given amplitude or absent from the T_2_ distribution. This constraint turns Eq.4 into a non-linear problem. For the solution to resemble the true T_2_ distribution, the T_2_ grid needs to contain a sufficient number and range of T_2*i*_ [16]. The T_2_ grid is established by logarithmic spacing between the lowest (T_2_,*min*) and highest T_2_, based on the number of allowed T_2_ times (nT_2_) and the expected width of the T_2_ distribution. More nT_2_ times may enhance confidence in the determination of the different components in the T_2_ distribution, but will prolong processing time. Most researchers utilize 40 nT_2_ times, encompassing T_2_ times between 15 ms and 2 s [46,60,61]. At these parameters, 30 min to several hours are required to compute a whole brain MWF map, depending on available hardware. We tested NNLS with nT_2_ of 20, 40, 80 and 120 T_2_ times, while varying the lower bound of the T_2_ grid, T_2,*min*_, between 3 – 15 ms in 1 ms intervals.

Assuming that 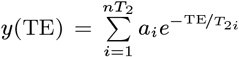, the signal S_*j*_ of echo j is defined by a system of linear equations

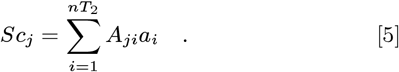

The matrix coefficients A_*ji*_ represent the exponentials, the decay signal computed by EPG. In principle, the amplitudes ai can be calculated by taking the pseudo-inverse of A_*ji*_. However, exact inverse solutions are contaminated by noisy data and no constraints, such as non-negativity of all ai, can be imposed. By contrast, NNLS allows for a misfit between the simulated and measured decay data yj, for instance due to noise, using the *χ*^2^ statistic (Eq.2). When minimizing

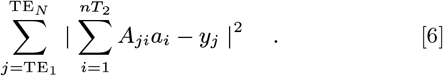

all ai are implicitly constrained to be non-negative by NNLS operating as an active set algorithm. The active set contains the regression coefficients, i.e. signal amplitudes, that are negative or zero when estimated without constraints. If the true active set were known, the solution would equal the unconstrained solution of all non-negative coefficients [62]. Starting with a zero vector where all coefficients are non-negative, NNLS approaches a prediction of the active set by iterative combination of all T_2,*i*_ with different amplitudes, fulfilling the non-negativity constraint.

Regularization is employed to produce continuous distributions of T_2_ relaxation times, which are expected to conform to reality. Thereby, adjacent peaks are merged to form a few distinct water pools. To constrain T_2_ distributions, an l2-norm regularization term is incorporated into Eq.6,

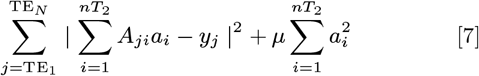

which minimizes the system’s “energy” to produce smooth, continuous T_2_ distributions [42]. By increasing the strength of the regularization, the central IEW peak becomes broader, while the short fraction size reduces. Other regularization, e.g. by the first or second order derivatives of the spectrum, may also be used [16,63].

Without regularization, 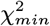 reflects the misfit in each data point by about one standard deviation [16]. With regularization, 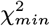 will be adjusted incrementally to increase *χ*^2^, since it is typically too low, fitting data too accurately [16,42]. The ratio of *χ*^2^ over 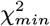 has commonly been set to 1.02 [60]. The misfit depends on the acquisition type and noise level.

To test the robustness of NNLS with respect to noise and the influence of regularization, both the decay curve SNR and the 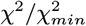 ratio were varied. White Gaussian noise was independently added to each of the computed decay curves (Matlab’s awgn function). Thereby, the power of the input signal was assessed before noise was added, yielding prescribed decay curve SNR levels of 30, 50, 100, 150, 200, 250, 300, 500 and 1000. The noisy data was in some cases compared to decay curves without added noise, i.e. no specific SNR level was generated. The regularization parameter was varied to yield 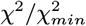 of 1.02, 1.05, 1.1, 1.15, 1.25, i.e. allowing misfits of 2 – 25%.

The estimated geometric mean (gm) T_2_ of the IEW and MW peak and the obtained MWF were compared to the ground truth to assess the accuracy of the NNLS solutions with EPG.

### In vivo data

To relate the simulations to in vivo imaging, MWI data from one female healthy volunteer (age 23 years) was collected. The volunteer gave written informed consent. One 32 echo GraSE T_2_ data set was acquired at a 3T Philips Achieva (Best, The Netherlands) with an eight-channel head coil, using TE/ΔTE/TR=10/10/1000ms [57]. The data were processed with the regularized NNLS, as described for the simulated data, using T_2,*min*_ = 15 ms, nT_2_ = 40 and 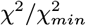 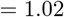, typical for in vivo multi-component T_2_ analysis [57,58].

## Results

### Extended Phase Graph – Flip Angle estimation

The reliability of the FA estimation by EPG depends predominantly on SNR. Figure 2 compares the FA estimates for the different acquisition types and decay curve noise levels. Despite greater variation from the true FA at lower SNR, estimated and true FAs still correlated linearly at all SNR levels (p < 0.023, r > 0.873). In the 32-echo simulation, the average slope was 0.836 ± 0.140 for all SNR levels. For 48-echoes and 64-echoes, FA estimates improved and the average slopes of the regression lines were 0.996 ± 0.060 and 1.022 ± 0.066, respectively. Note that EPG estimates B1^+^ inhomogeneities relative to the assumed refocusing angle of 180°, so that larger FAs, e.g. 190°, are displayed as 170°.

**Fig. 2.**
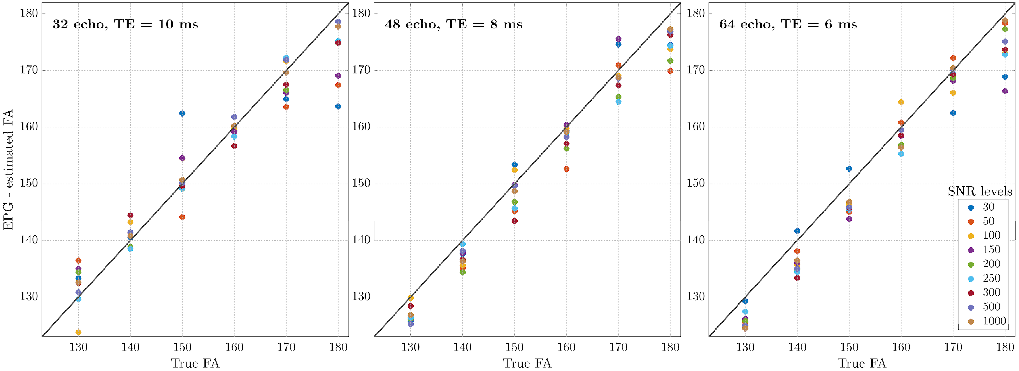
EPG-FA estimation in the presence of stimulated echoes and at different noise levels in the highest myelin water model (MWF = 21%). Independent of the acquisition type and for most SNR levels, EPG estimated FAs were within a few degrees of the true FA value, indicated by the black line. At very low SNR levels, deviations between the true and estimated FA were noticeable, but lower and higher FAs could still be well distinguished.

Table 1 summarizes the SNR-dependent root-mean squared error (RMSE) of the FA estimates across different acquisition types and the three MWF models (M1, M2, M3). The three acquisition types are referred to by their ETL: 32e, 48e and 64e indicating acquisitions of 32 echoes with 10 ms ΔTE; 48 echoes with 8 ms ΔTE and 64 echoes with 6 ms ΔTE, respectively. With improving decay curve SNR, FA estimates became more reliable. Similar RMSE estimates indicated little difference between acquisitions using different ΔTE. Across all noise levels, models and acquisition types, the estimated FAs were 127.8 ± 3.6°, 138.4 ± 3.7°, 148.6 ± 3.9°, 158.9 ± 3.2°, 168.5 ± 3.1°, 173.5 ± 4.2° relative to the true FAs of 130°, 140°, 150°, 160°, 170° and 180°. The apparent difficulty in capturing FA = 180° (see also Fig. 2) is due to the FA limit of 180° with EPG. Small systematic deviations from 180°, however, are of little consequence for the EPG signal decay.

Given that only a limited number of T_2i_ times is used to describe the T_2_ decay, some variance is expected with respect to nT_2_ and the choice of T_2,*min*_. For a particular acquisition type and model (M2, 48e, MW T_2_ =10 ms, SNR = 500), changing nT_2_ yielded on average variations of less than 1° across all FAs. Changing the MW T_2_ led to arbitrary, nonsystematic variations in the FA estimation. On the other hand, changing T_2,*min*_ from the lowest to highest (3 ms to 15 ms) resulted in variations of 1° – 4° from the true FA with a tendency towards higher FAs for longer T_2,*min*_. Regularization played no role in the FA estimation.

### Decay curve SNR and B1^+^ inhomogeneities: Comparison of simulated and in vivo data

The SNR of the decay curves affects the FA estimation, with low SNR also resulting in significant residuals. Importantly, while lower FAs produce systematically lower signal at the first echo by inefficient re-focusing, a signal increase due to positive noise at the first echo could lead to an overestimation of the MWF. Together B1+ inhomogeneity and the decay curve SNR contribute to the estimated SNR (SNRest) that is obtained from the NNLS fit.

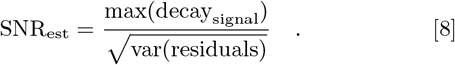

The surface plot in Figure 3A exemplifies the influence of these two parameters on SNRest for M2, 32e. In vivo data acquired at 3 T are depicted for comparison in panel B.

**Fig. 3.**
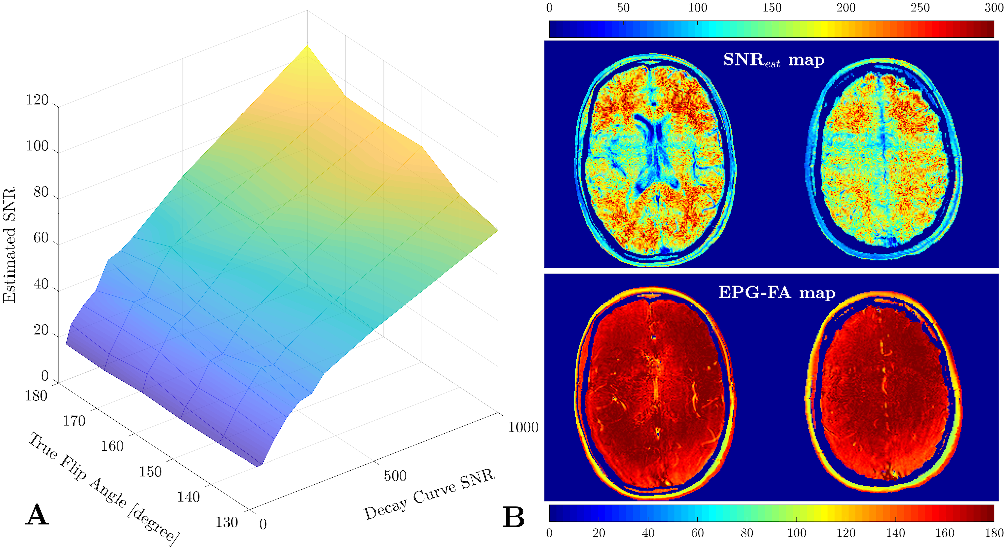
A) Estimated SNR, given the maximum signal of the decay curve and the residuals, relative to the true FA and SNR that was assigned to the decay. As expected, larger initial SNRs and larger FAs yielded larger SNR_*est*_. (B) In vivo SNRest and EPG-FA map are displayed for two different brain slices. SNRest is indirectly dominated by the local FA via the steady-state + receive sensitivity of the 8-channel head coil. A direct spatial correlation with the EPG-FA maps, however, cannot be seen. SNRest values clearly distinguished white and gray matter regions, with WM SNRest being > 180.

In vivo data yielded SNRest values > 180 and FAs of > 150 for most WM. By comparison, SNRest of the simulation at FAs around 180° and signal SNR = 1000 was just over 100. Thus, particularly in WM, the decay curve SNR at respective FAs in vivo appears to be sufficient for T_2_ estimation, and the simulated data represent possibly worse signal SNR scenarios than encountered in vivo at 3 T. This is further supported by the direct comparison of the decay curves (Figure 4). At comparable FAs, the decay curves displayed similar stimulated echo and noise patterns for high SNR values. Thus, all subsequent figures and computations will evaluate the decay curves with a simulated decay curve SNR = 1000.

**Fig. 4.**
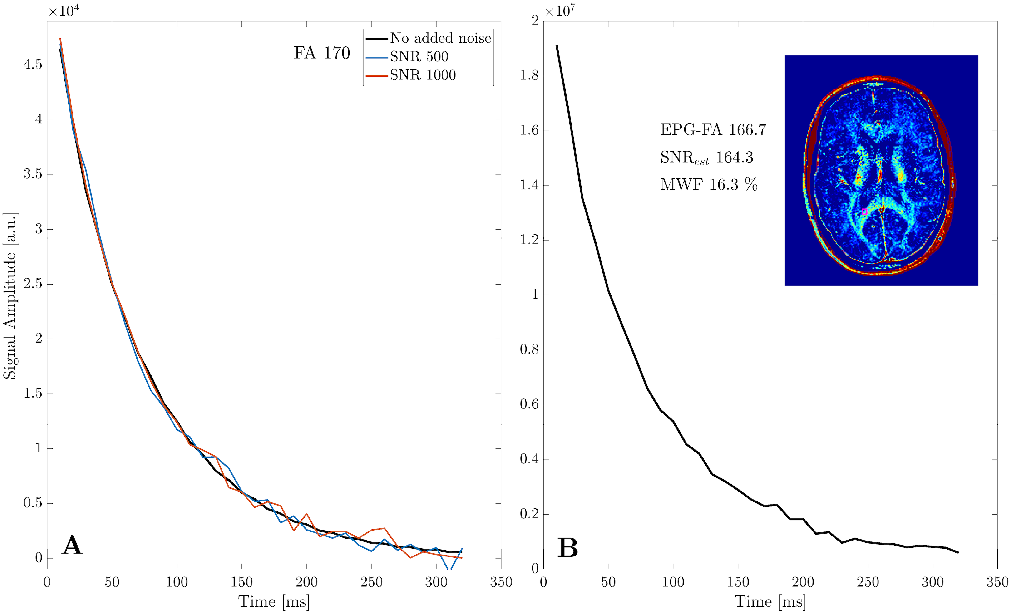
Direct comparison of simulated (A) and experimental (B) decay curves for a typical WM voxel, here of the splenium of the corpus callosum, yielding FA and MWF estimates in line with the simulation (M2, 32e, FA 170^°^). Note that ‘no added noise’ means no white Gaussian noise was added to the decay curve, but Gaussian noise was present in the T_1_ and T_2_ distributions of the simulated image voxel.

### Relationship between T_2,*min*_ and MW T_2_ time

To understand the relationship between the choice of T_2,*min*_ and the MW T_2_ time, it is most informative to look at the regularized T_2_ distributions (Figure 5). Independent of T_2,*min*_, FA, noise level or MW T_2_, the IEW peak was centered at the true IEW T_2_ time. Nonetheless, the gmT_2_ deviated slightly from the true value (IEW gmT_2_ time: panel A 66.4 – 67.5 ms, B 69.3 – 70.2 ms, C 71.0 – 71.4 ms), likely introduced by the limited data sampling of the T_2_ distributions. For instance, with nT_2_ = 40, 4 – 7 data points characterized the IEW peak. Choosing larger nT _2_may provide more stable estimates of the IEW gmT_2_. But, the gain in better estimating the IEW gmT_2_, relative to the choice of nT_2_, is small (about 1*ms*) and computational time must be weighed against minimally improved accuracy.

**Fig. 5.**
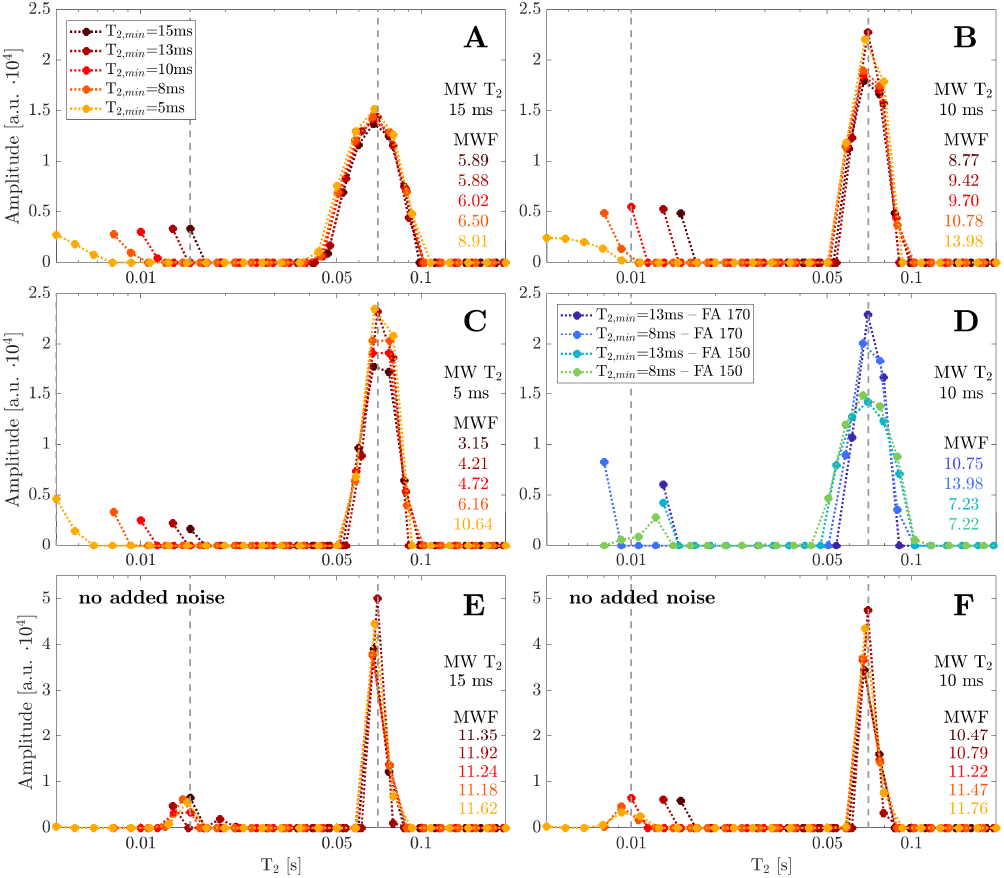
Comparison of T2 distributions obtained by regularized NNLS with varying T_2,*min*_, noise levels and FAs. Data shown here correspond to M2, i.e. 11.6% MWF, 48e and nT_2_ = 40. Panels A – D correspond to decay curves with SNR = 1000 compared to E and F, where no additional noise was applied to the decay curves. All red-color scheme curves (A – C, E & F) correspond to FA = 180° and D shows examples of lower FAs. Vertical dashed lines indicate the true MW T_2_ and IEW T_2_ times. The 5 ms MW T_2_ line in panel C is co-localized with the y-axis. IEW T_2_ was 70 ms in all simulations. The MW T_2_ times are listed on the right side of each plot. The MWF estimate for each T_2,*min*_ is quoted in the lower right hand corner of each panel.

Due to its rapid decay and lesser signal, the MW compartment is more sensitive to noise than the IEW component. Hence, the MW pool was typically represented by only one or two elevated points rising towards T_2,*min*_, rather than a peak (panels A – D), indicating that our acquisitions and T_2_ distributions have insufficient range to fully describe the MW signal. On the contrary, T_2_ distributions representing decay curves with no added Gaussian noise (E & F), explicitly captured the influence of T_2,*min*_ on the MW peak. In panel F, when T_2,*min*_ > MW T_2_, the MW “peak” was a slope, characterized by a single T_2*i*_ time, pushing towards the lowest allowed T_2_, i.e. T_2,*min*_. Once T_2,*min*_ < MW T_2_, small peaks formed that fully depicted the short T_2_ signal and the MW gmT_2_ resembled the true MW T_2_. For MW T_2_ = 10 ms (panel F), the estimated MW gmT_2_ was 15, 13, 10, 9.5, 9.4 ms for T_2,*min*_ of 15, 13, 10, 8 and 5 ms. In the presence of added noise (panels A – D), both T_2_ peaks became wider, which made it particularly difficult to capture the short T_2_ component fully.

It needs to be determined how strongly the MWF assessment depends on the selection of T_2,*min*_ and the estimation of the MW gmT_2_. Furthermore, it needs to be ascertained how short T_2,*min*_ can be selected relative to the shortest acquisition TE (e.g. 6, 8 or 10 ms) to recover the true MWF.

Estimating the geometric mean T_2_ of the intermediate water – IEW – compartment In healthy WM, the intermediate T_2_ is assigned to water in the axonal lumen, cell bodies and interstitium. The IEW peak is relatively broad, with a gmT_2_ of approximately 68 – 72 ms at 3 T [59]. Therefore, varying nT_2_ hardly influenced the determination of IEW gmT_2_ (variations approximately 1 ms). Increasing regularization progressively reduced the IEW gmT_2_ from the true time of 70 ms. Lower FAs did not systematically underestimate IEW gmT_2_. Differences in T_2,*min*_ changed IEW gmT_2_ by less than 3 ms. When the IEW gmT_2_ was not stable with respect to T_2,*min*_, it approached the true value when the T_2,*min*_ was closer to the MW T_2_ (Supplementary Table 1). If T_2,*min*_ was longer than the MW T_2_, IEW gmT_2_ increased above the true value, in line with the observations from Fig. 5C. In turn, for long MW T_2_ (20 ms, Fig.5A), the IEW gmT_2_ was underestimated. The amount of MW did not affect the IEW gmT_2_ estimation.

Across all FAs, T_2,*min*_ and MW T_2_ times, we obtained 66.2 ± 2.2 ms, 66.7 ± 2.1 ms and 66.4 ± 2.2 ms for M1 (32/48/64e,respectively), 66.7 ± 1.6 ms, 66.8 ± 1.6 ms and 66.8 ± 1.6 ms for M2, and 67.6 ± 1.5 ms, 67.7 ± 1.6 and 67.6 ± 1.5 ms for M3, yielding an approximate variation of 3 – 4 ms from the true value 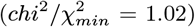. Without the addition of noise to the decay data, IEW gmT_2_ times were closer to the ground truth (less than 2 ms difference) and demonstrated a dependency on FA. For M1, 32e the obtained IEW gmT_2_ was 69.5±0.8 ms at 180°, 69.2 ± 0.9 ms at 170°, 68.4 ± 1.4 ms at 150° and 68.4 ± 1.5 ms at 130°.

### Estimating the geometric mean T_2_ of the MW peak and the MWF

When comparing the true and estimated MW gmT_2_ at different FAs (M2 - 48e), the three MW T_2_’s of 20, 13 and 8 ms were only accurately reproduced at relatively high FAs and T_2,*min*_ < MW T_2_. Once T_2,*min*_ < MW T_2_, the MW gmT_2_ reached a plateau and the solution remained stable in most cases (Figure 6). However, the plateau gmT_2_ time was typically shorter than the true MW T_2_, particularly at lower FAs. If TT_2,*min*_ was very short, MW gmT_2_ tended to decline further. Moreover, it can be difficult to reach a stable solution, if MW T_2_ is short, e.g. MW T_2_ = 8 ms. In the absence of added noise (lower right panel) a similarly plateauing behavior was observed, with some plateau values closer to the true T_2_ time than in the SNR = 1000 case. Again, the dependence on FA was less pronounced when noise was added to the decay curves. Without added noise, the median MW gmT_2_ for the 20 ms MW T_2_ was 19.5, 19.4, 17.5 and 14.6 ms at 180°, 170°, 160°, 150°. For 13 ms MW T_2_, we obtained 12.7, 12.9, 11.8 and 11.7 ms at the decreasing FAs and 9, 9, 9 and 9 ms relative to the 8 ms true MW T_2_ for the same M2, 48e acquisition.

**Fig. 6.**
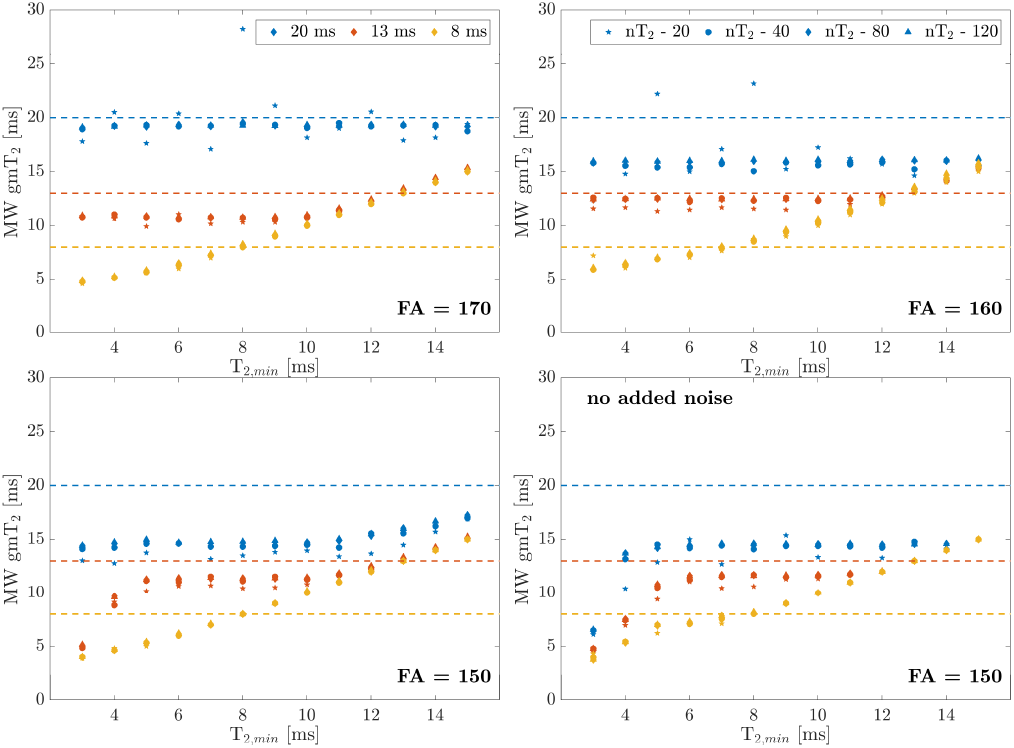
MW gmT_2_ estimates for various FAs, T_2,*min*_, nT_2_s and true MW T_2_ times. Typically, MW gmT_2_ estimates yielded stable solutions if T_2,*min*_ ± MW T_2_, however, the stable solution may estimate gmT_2_ to be shorter than the true MW T_2_. Here, the SNR was 1000 for all panels except the bottom right panel.

Variations with nT_2_ were on average below 1 ms, with nT_2_ = 20 T_2_ times yielding the largest offsets from the other nT_2_ estimates. Larger nT_2_ tended to increase the MW gmT_2_ minimally. Stronger regularization led in some cases to much longer MW gmT_2_, presumably a result of coalescence of the two water peaks. Correspondingly, with increasing regularization, the MWF tended to decrease initially, and then increase as the peaks merged and the consolidated peak extended into the MW T_2_ window. Table 2 compares the estimated MW gmT_2_ time (left) in the three models and for the three different acquisition approaches relative to the selected T_2,*min*_, and contrasts gmT_2_ estimates with the obtained MWF values (right).

MW gmT_2_ estimates improved in many cases for acquisitions with shorter echo spacing. Particularly the 48e, 8 ms echo spacing acquisition showed stable estimates, close to the true MW T_2_ time of 20 ms. Notably, even in the presence of unstable MW gmT_2_ estimates, MWFs were consistent across T_2,*min*_, albeit underestimated for the 20 ms MW T_2_. Figure 7 summarizes the relationship between MWF, MW T_2_ and selection of T_2,*min*_. Data for 48e and 64e are shown; 32e data are provided in Supplementary Figure 1. The top row displays the mean behavior in estimated MWF across MW T_2_ times of 15, 13, 10 and 8 ms relative to T_2,*min*_, since typically the true T_2_ time of the MW compartment is unknown. The bottom row illustrates at the relationship of MWF and individual MW T_2_ after selection of T_2,*min*_.

**Fig. 7.**
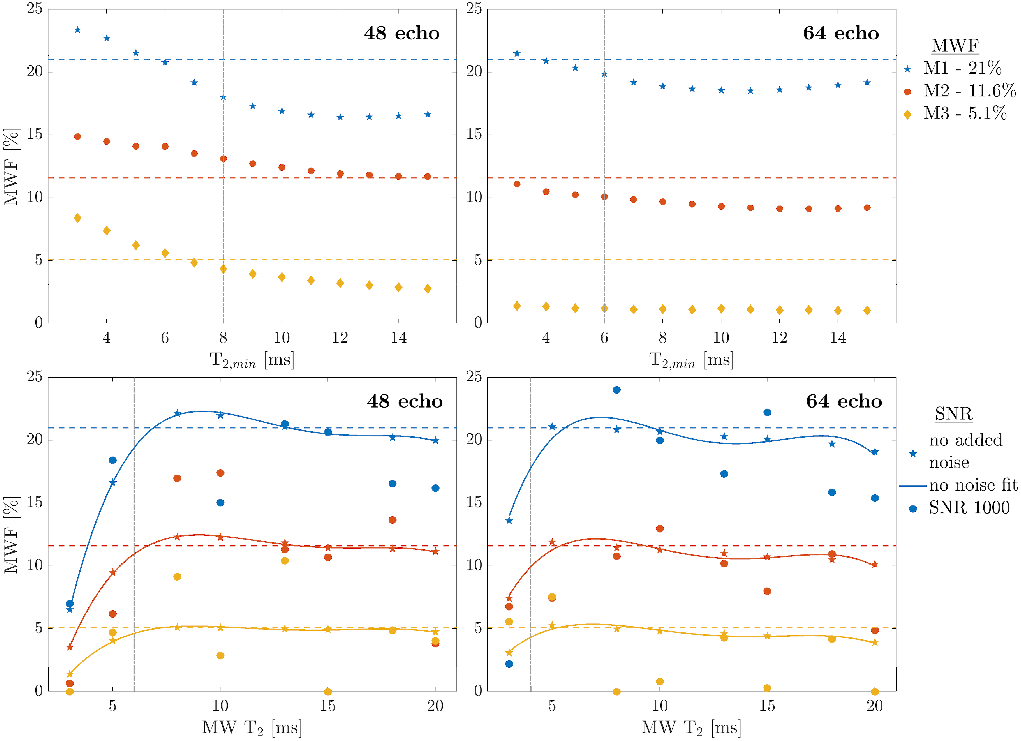
Mean MWF estimates relative to the true values at different T_2,*min*_ (top) and MW T_2_ times (bottom), at SNR = 1000, *χ*^2^ = 1.02, nT_2_ = 120. Top row: Mean MWF averaged over different MW T_2_ times (15, 13, 10 and 8 ms) for different acquisition strategies at FA = 170°. The true MWF values (21, 11.6 and 5.1%) are shown for comparison as dashed lines. The gray, dash-dotted verticals indicate the acquisition TE, relative to T_2,*min*_. On average, MWF values are closest to the true value when T_2,*min*_ is chosen to be just below TE1. Bottom row: Mean MWF values for different MW T_2_ times computed at a specified T_2,*min*_ (gray, dash-dotted vertical line). Estimates are compared in the presence (filled circles) and absence of noise (filled stars). A fourth-order polynomial fit was applied to the no added noise data. Even in the absence of noise, the MWF is strongly underestimated if MW T_2_ is shorter than T_2,*min*_ = TE – 2 ms. For all MW T_2_ that are longer than the selected T_2,min_ time, the estimated MWF approaches the true value. At SNR 1000 (filled circles), estimated MWF vary more around the true value. Note that the x-axes differ; top row T_2,*min*_, bottom row MW T_2_.

Average MWF estimates were relatively stable for T_2,*min*_ > TE1, although often underestimated (top row). In most cases, the estimated MWF was closest to the true value if T_2,*min*_ was just below or on the order of TE1. However, the exact relationship depended on the true MW T_2_. Selecting T_2,*min*_ according to TE1, i.e. 8 ms for 10 ms TE, 6 ms for 8 ms TE, and 4 ms for 6 ms TE, we obtained 14.63 ± 6.61%, 17.66 ± 5.57% and 17.79 ± 7.28% for 32e, 48e and 64e, across all possible MW T_2_ times (bottom row, filled circles). These estimates were lower than the true MWF of 21% due to the inclusion of MW T_2_ times shorter than T_2,*min*_. Excluding data for which MW T_2_ < T_2,*min*_, the average MWFs were 17.00 ± 5.82%, 19.31 ± 4.17% and 20.01 ± 3.95% for 32e, 48e and 64e, respectively, for M1, 12.19 ± 5.46%, 12.32 ± 5.00% and 9.32 ± 2.71% for M2 (true fraction 11.6%) and 3.76 ± 3.07%, 5.23 ± 3.92% and 2.45 ± 2.94% for M3 (true fraction 5.1%), at appropriately chosen T_2,*min*_. Shorter echo spacing improved the consistency of the MWF estimation across MW T_2_ times, as reflected in the smaller standard deviations. Regardless, MWFs differed insignificantly between acquisition schemes, if T_2,*min*_ was chosen as described. Overall, the true MWF was underestimated by 0.13-4 percentage points (at 170°, SNR = 1000, 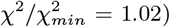). In the absence of added noise (filled stars and fitted line), MWF values demonstrated an even higher degree of similarity, deviating from the true value by 0.02 - 0.64 percentage points. Specifically, we obtained 20.36 ± 0.31% for 32e, 21.02 ± 0.90% for 48e and 20.26 ± 0.69% for 64e, close to the true value of 21% MWF. For M2, we found 11.23 0 ±.24%, 11.73 ± 0.49% and 11.00 ± 0.60% and for M3 4.62 0 ±.17%, 4.97 0 ±.14% and 4.62 ± 0.46%, respectively, for 32e, 48e and 64e at appropriately chosen T_2,*min*_.

## Discussion

### Importance of B1^+^ homogeneity and SNR

The EPG algorithm has much enhanced our ability to accurately interpret T_2_ decay data in the presence of stimulated echoes [45]. Yet, even when modeling spurious echoes in the T_2_ decay, imperfect refocusing pulses yield reduced signal amplitudes at early odd echoes, which leads to slight, systematic underestimations in gmT_2_ time and MWF. Notably, underestimations in MWF and IEW gmT_2_ also occur in the presence of noise44. Thus, while the influence of sub-optimal refocusing FAs and increased noise are clearly characterized individually, their combined presence leads to less systematic, and therefore less interpretable effects. Although noise affects predominately the residuals of the later echoes, noise in earlier data points may be mistakenly interpreted as stimulated echo artifact and vice versa.

Here, we used small-scale constraints, which increased *χ*^2^ to raise accuracy in the NNLS component estimation. Graham et al. showed that least squares-based constraints, as compared to *χ*^2^, may be advantageous at intermediate to low SNR in obtaining acceptable accuracy in determining multiple T_2_ components [42]. In early years of multi-echo T_2_ analysis, acquisition parameters of TE = 10 ms, 32 echoes and voxel SNR of 100, as determined at TE = 0 ms and the standard deviation of the noise, did not match the optimal TE of 6.4 ms and SNR of 700, then quoted to be required to resolve multiple T_2_ components by NNLS [42]. As shown in Figure 3, these limits may now be regarded as too strict. With advancing MR hardware and newer acquisitions, we are coming closer to achieving optimal data collection, and traditional restrictions for multi-component T_2_ analyses may need to be revised. Nevertheless, regularization should be used conservatively, allowing data misfit of only a few percent, typically 1 or 2 % [64] of that of the unregularized *χ*^2^, as stronger Tikhonov regularization quickly affects both MWF and IEW T_2_ measurements. Increasing noise by itself will also lead to widening of the T_2_ peaks (Fig.5), reflecting the uncertainty in the T_2_ estimation.

Regardless of the noise level, EPG captured the true FAs on average within an accuracy of 1 – 4°. Errors tended to be larger at lower FAs and lower decay curve SNRs. FAs at 180° appeared to be less accurately estimated. This is in part due to the EPG estimation limit of FAs < 180°, so that FAs will always be underestimated at this limit. Moreover, the trigonometric relationship between signal loss and FA plateaus at 180°, making the identification of the correct FA more challenging. Stimulated artifacts are, however, very small at FAs between 165° and 180°.

**Table 2.**
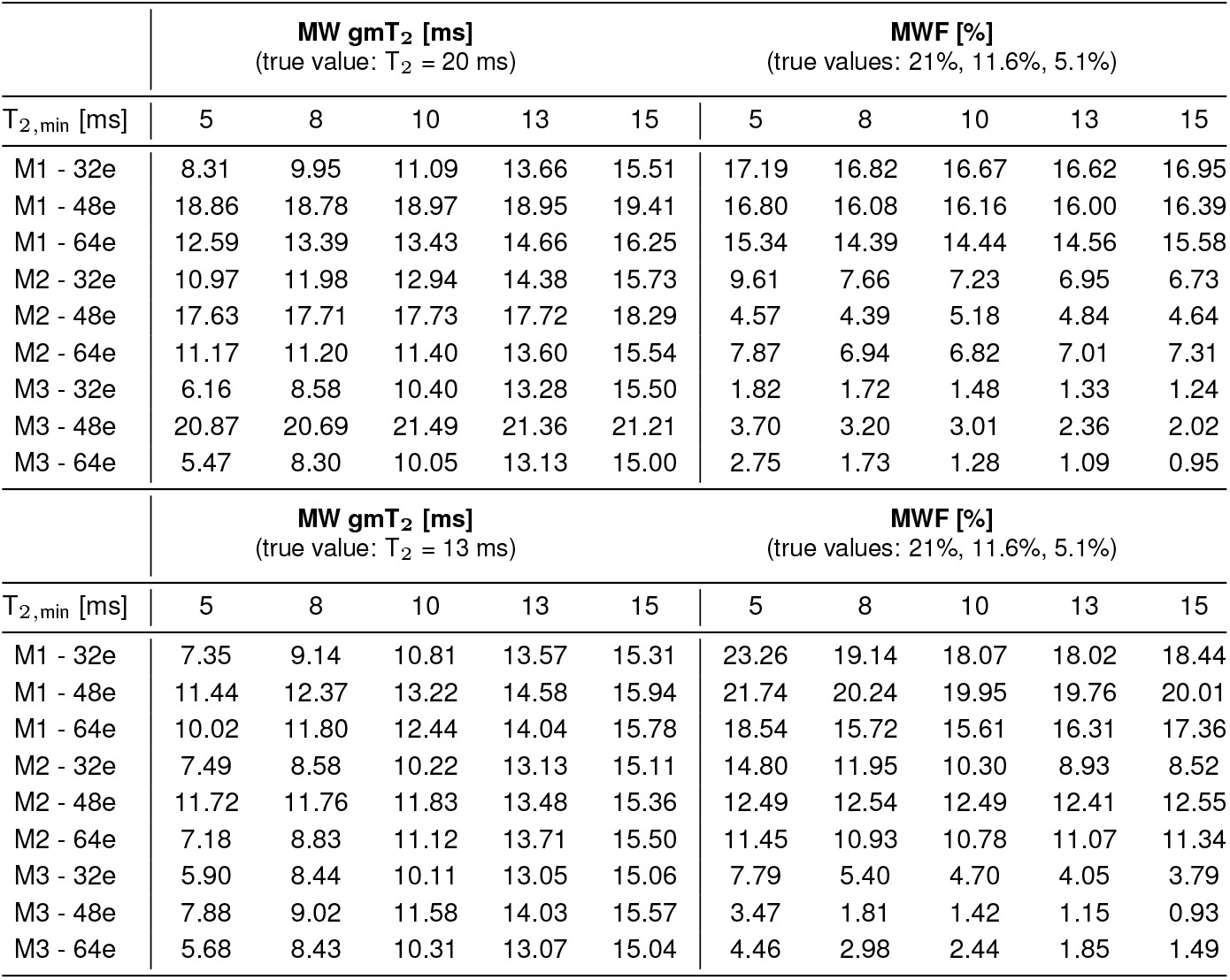
Influence of the choice of acquisition and selection of T_2,min_ on accurate estimation of the MW T_2_ and MWF. The top part of the table evaluates models with true MW T_2_ of 20 ms, and the bottom half with MW T_2_ of 13 ms. Values shown correspond to 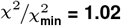 nT_2_ = 120, SNR 1000, averaged over FAs of 150° â€” 180°. True MWF values are model dependent Model 1 (21% MWF) / Model 2 (11.6%) / Model 3 (5%).

|  | MW gmT<sub>2</sub> [ms]<br>(true value: T<sub>2</sub> = 20 ms) |  |  |  |  | MWF [%]<br>(true values: 21%, 11.6%, 5.1%) |  |  |  |  |
| --- | --- | --- | --- | --- | --- | --- | --- | --- | --- | --- |
| T<sub>2,min</sub> [ms] | 5 | 8 | 10 | 13 | 15 | 5 | 8 | 10 | 13 | 15 |
| M1 - 32e | 8.31 | 9.95 | 11.09 | 13.66 | 15.51 | 17.19 | 16.82 | 16.67 | 16.62 | 16.95 |
| M1 - 48e | 18.86 | 18.78 | 18.97 | 18.95 | 19.41 | 16.80 | 16.08 | 16.16 | 16.00 | 16.39 |
| M1 - 64e | 12.59 | 13.39 | 13.43 | 14.66 | 16.25 | 15.34 | 14.39 | 14.44 | 14.56 | 15.58 |
| M2 - 32e | 10.97 | 11.98 | 12.94 | 14.38 | 15.73 | 9.61 | 7.66 | 7.23 | 6.95 | 6.73 |
| M2 - 48e | 17.63 | 17.71 | 17.73 | 17.72 | 18.29 | 4.57 | 4.39 | 5.18 | 4.84 | 4.64 |
| M2 - 64e | 11.17 | 11.20 | 11.40 | 13.60 | 15.54 | 7.87 | 6.94 | 6.82 | 7.01 | 7.31 |
| M3 - 32e | 6.16 | 8.58 | 10.40 | 13.28 | 15.50 | 1.82 | 1.72 | 1.48 | 1.33 | 1.24 |
| M3 - 48e | 20.87 | 20.69 | 21.49 | 21.36 | 21.21 | 3.70 | 3.20 | 3.01 | 2.36 | 2.02 |
| M3 - 64e | 5.47 | 8.30 | 10.05 | 13.13 | 15.00 | 2.75 | 1.73 | 1.28 | 1.09 | 0.95 |

|  | MW gmT<sub>2</sub> [ms]<br>(true value: T<sub>2</sub> = 13 ms) |  |  |  |  | MWF [%]<br>(true values: 21%, 11.6%, 5.1%) |  |  |  |  |
| --- | --- | --- | --- | --- | --- | --- | --- | --- | --- | --- |
| T<sub>2,min</sub> [ms] | 5 | 8 | 10 | 13 | 15 | 5 | 8 | 10 | 13 | 15 |
| M1 - 32e | 7.35 | 9.14 | 10.81 | 13.57 | 15.31 | 23.26 | 19.14 | 18.07 | 18.02 | 18.44 |
| M1 - 48e | 11.44 | 12.37 | 13.22 | 14.58 | 15.94 | 21.74 | 20.24 | 19.95 | 19.76 | 20.01 |
| M1 - 64e | 10.02 | 11.80 | 12.44 | 14.04 | 15.78 | 18.54 | 15.72 | 15.61 | 16.31 | 17.36 |
| M2 - 32e | 7.49 | 8.58 | 10.22 | 13.13 | 15.11 | 14.80 | 11.95 | 10.30 | 8.93 | 8.52 |
| M2 - 48e | 11.72 | 11.76 | 11.83 | 13.48 | 15.36 | 12.49 | 12.54 | 12.49 | 12.41 | 12.55 |
| M2 - 64e | 7.18 | 8.83 | 11.12 | 13.71 | 15.50 | 11.45 | 10.93 | 10.78 | 11.07 | 11.34 |
| M3 - 32e | 5.90 | 8.44 | 10.11 | 13.05 | 15.06 | 7.79 | 5.40 | 4.70 | 4.05 | 3.79 |
| M3 - 48e | 7.88 | 9.02 | 11.58 | 14.03 | 15.57 | 3.47 | 1.81 | 1.42 | 1.15 | 0.93 |
| M3 - 64e | 5.68 | 8.43 | 10.31 | 13.07 | 15.04 | 4.46 | 2.98 | 2.44 | 1.85 | 1.49 |

Although it is important to compute the actual FAs and characterize stimulated echo artifacts in the T_2_ decay curves, EPG adds to the computational post-processing time. Replacing the FA estimation by EPG with acquired FA maps could lessen the required processing time. FA maps are typically acquired at a lower spatial resolution than multi-echo T_2_ data, as B1^+^ is assumed to be spatially slowly varying. Using spatial constraints on T_2_ distributions and FAs across anatomical and pathological structures, as recently described [65], has the potential to improve noise robustness for multi-component T_2_-mapping.

We want to emphasize again that EPG does not reverse signal loss due to low FAs or the resulting underestimation of the MWF. Thus, improving B1^+^ homogeneity, such that refocusing FAs are near 180°, should be considered a priority. Greater B1^+^ homogeneity can be achieved by dielectric pads or parallel excitation by multi-element transceiver arrays. At 3 T, underestimations in MWF may be considered less significant than at higher field strengths, where greater B1^+^ inhomogeneity is likely to impose larger spatial variations in MWF. However, as the combination of FA and noise, i.e. the SNRest, will be the determining factor for the MWF estimation, systematic underestimations in MWF measurements may be alleviated as increased SNR may make up for greater B1^+^ variations at higher field strength. Even more, SNR profiles of multi-channel receive coils, which provide greatest SNR near the edge of the head, oppose the typical B1^+^ profile. Thus, these coils may avoid the discussed underestimations. Further work is needed to explore B1^+^ penalties as well as acquisition strategies at 7 T that can appropriately capture the shorter T_2_ decay of MW within specific absorption rate (SAR) limits.

### Regularization

Although regularization is useful for resolving multi-components of the T_2_ distribution in the presence of noise, it comes at the cost of underestimating the MWF. We also noted that the IEW gmT_2_ time shortened with increasing regularization, albeit underestimations were less than 1 ms from the true value44. *χ*^2^ constraints should only be applied over a narrow range that ensures accurate detection of multiple T_2_ components [42]. Increasing the “energy” constraint, i.e. enforcing greater smoothing by Tikhonov regularization, widens and subsequently merges different water peaks, leading to shorter IEW gmT_2_ and reduced apparent MWF. As the coalesced peak becomes even wider, it can partly extend into the a priori defined short T_2_ interval, increasing measured MWF. Other regularization approaches have focused on data denoising [66] or used spatial constraints for handling of noisy data [65,67–69]. Although these methodologies yield visually appealing results, the impact of regularization should be carefully evaluated. As discussed, noise cannot always be distinguished from decay curve alterations due to stimulated echoes. Moreover, voxels of relatively low myelin content may appear noisy and residual myelin signal may be falsely removed. Spatial regularization may also have limited success for smaller tracts, where partial volume related underestimations in the MWF may propagate into the region-of-interest.

### Other methodologies for mapping of myelin water

Different myelin water imaging techniques have been reviewed by Alonso-Ortiz et al.[70]. Multi-echo gradient-echo imaging has gained popularity for multi-compartmental fitting of T_2_* decay data [71,72]. Despite the simpler and faster acquisition and lower SAR, increased sensitivity to static field inhomogeneities [73] and physiological noise [74] have made it challenging to robustly obtain T_2_* myelin maps comparable to those obtained using spin-echo based techniques [71,75]. Importantly, while NNLS17 enables determination of an unlimited and unknown number of T_2_ components [16], most T_2_*-based techniques assume a fixed number of water compartments, usually two or three [72,76–78]. Currently, no NNLS algorithm exists that can separate the T_2_* magnitude and frequency offsets that need to be modelled for an unknown number of signal components. A priori imposition of the number of water pools may hamper the use of these techniques and other model-based approaches [79] in pathological cases, where changes in T_2_ and additional water pools have been reported [23,36,80]. In this work, only the multi-component T_2_ acquisition and NNLS analysis were considered, although other acquisition types for myelin water imaging have been proposed [81–83] and different analysis techniques may be employed [84,85]. To date, most of these approaches lack thorough histopathological validation and some are known to depend on the presence of magnetization exchange [86,87].

### T_2,*min*_, MW T_2_ and MWF

Using NNLS, MW T_2_ may be inaccurately estimated in the presence of noise as the MW gmT_2_ computation depends on T_2,*min*_ and acquisition echo spacing. Typically, MW gmT_2_ will reach a stable solution if T_2,*min*_ ≤ MW T_2_, i.e. when the short T_2_ peak is fully captured. Note that the MW T_2_ is currently not regarded as having clinical relevance. Our observation of stable MWF even in the presence of changing MW gmT_2_ (Tab. 2) further supports the use of MWF over MW gmT_2_. All dependencies are summarized in Table 3.

We observed that the MW peak was best captured if T_2,*min*_ was somewhat shorter than TE1 and shorter or equal to the MW T_2_ in order to reproduce the true MWF. Choosing T_2,*min*_ much lower than TE1 resulted in artifactually high MWF values due to noise in the first data points. Interestingly, longer MW T_2_ time did not necessarily result in more accurate gmT_2_ and MWF values, despite both T_2,*min*_ and the acquisition TE1 being sufficiently short to capture the MW signal. Previous work suggested that longer MW T_2_ may require a longer first TE to obtain optimal SNR for the description of the water pools [41]. This is in line with our observations, suggesting that the optimal TE for MWI may need to be close to the MW T_2_.

Hardware restrictions make it difficult to accurately determine very short T_2_ times as echo spacing is limited in current acquisitions. Because the in vivo MW T_2_ is unknown, further work is needed to determine the relevance of ΔTE in accurately estimating the MWF. The increased number of echoes necessary with shorter ΔTE will require longer TR to comply with SAR restrictions, possible posing a limit on measurement time. Undersampling approaches for spatial encoding, such as compressed sensing [88,89], may help to offset such limits.

Finally, one fixed set of T_2_ boundaries and parameters is typically chosen, which is believed to characterize the IEW and MW pools on average. Note that we did not discuss the T_2_ cut-off time here that separates thesewater pools. This parameter was well established in our simulations, but typically needs to be determined from the T_2_ distributions. As shown, single voxel MWF computations in vivo are challenging and should be approached with caution as noise and natural variations in myelin across the WM affect the MWF computation. Voxel-based average MWF measures are most commonly performed. They provide better reproducibility than region-based analyses, which would yield only a single regional MWF [44,60]. Across different MW T_2_ times, in the presence of noise and due to imperfect refocusing FAs, we found that the true MWF can be estimated at 0.13 – 4 percentage point deviation from the true value. Our simulations suggest that in vivo MWF measurements are reproducible [90], albeit underestimated. Reducing T_2,*min*_ below the first TE appears to reasonably recover some signal lost due to regularization and FA imperfections. For comparison of data from different sites and different acquisition protocols, average SNRest and or FAs may need to be reported to assess MWF differences.

### Limitations

The translation of our observations to in vivo scenarios is skewed by our limited knowledge of the true T_2_ distribution. We attempted to cover a wide range of acquisition and analysis parameters, relevant to the estimation of MWF. We did not simulate non-equidistant refocusing pulse schemes, previously proposed to better capture long T_2_ components [43], and also limited our simulations to pure spin-echo T_2_ decays, rather than the gradient and spin-echo acquisition currently used [58]. Non-uniform echo spacing implies a non-linear increase in refocusing pathways, greatly prolonging the computation of all magnetization states with EPG [7]. It is thus not commonly used. Computational time for EPG remains manageable if the echo spacing is changed by multiples of TE. Only Gaussian noise was investigated, as other noise patterns, such as Rician noise, have been described to affect MWF measurements similarly [44]. We chose Gaussian noise, because it is predominantly present in the earlier echoes that capture the rapid signal decay of the MW compartment. The present Bloch simulation ignored effects such as diffusion or exchange [91,92]. In the presence of exchange, an MWF decrease may be associated with a decrease in the IEW gmT_2_ [93]. Such exchange-associated shift in IEW gmT_2_ would be larger than what we observed here by altering acquisition and post-processing parameters [94]. We did not cover scenarios with MWF close to zero, as might be the case in neonates, completely demyelinated MS plaques or gray matter [95]. The created MWF scenarios only differed in the number of myelin bilayers, but kept the thickness or spacing of individual sheaths unchanged. Such changes, however, might occur due to swelling or vacuolization. This simplified model geometry created realistic water pool fractions and permits incorporating the above described parameters in the future. Although the pool fractions were the determining factor in this work, less so the actual geometry and distribution of the myelinated axons, the chosen setup allowed us to compute field inhomogeneities [96] and assign different compartmental resonance frequencies, which have been described in the literature. However, without modeling of diffusion, central k-space echoes will only minimally be affected. Finally, our simulations allowed generation of decay curves with stimulated echo artifacts independent of the EPG, which was used to analyze the decay curves.

**Table 3.** Summary of the investigated outcome measures and their dependence on nT_2_, regularization and the relationship between MW T_2_, T_2,min_ and acquisition echo spacing ΔTE. Average differences from the true values are indicated for available data. NI: not investigated. pp: percentage points

| | $nT_2$ | $\chi^2/\chi_{min}^2$ | MW $T_2$ | $T_{2,min}$ | $TE_1$ and $\Delta TE$ |
| --- | --- | --- | --- | --- | --- |
| **FA estimation** | $\sim 1^\circ$ | | Non-systematic variations | NI | In some cases, improved with shorter $TE_1/\Delta TE$ |
| **IEW gm$T_2$** | $\sim 1$ ms | Decreasing with increasing $\chi^2$ | Overestimated if $T_{2,min} > MW T_2$<br>Underestimated if $T_{2,min} < MW T_2$ | $< 3ms$<br>Most accurate: $T_{2,min} \sim MW T_2$ | No difference |
| **MW gm$T_2$** | $< 1$ ms | Increasing with increasing $\chi^2$ | | Stable, but underestimated, if $T_{2,min} < MW T_2$ | In some cases, improved with shorter $TE_1/\Delta TE$ |
| **MWF** | NI | Decreasing initially, then increasing with $\chi^2$ | Stable if MW $T_2 > TE$<br>Remains stable if MW $T_2 > (TE-2ms)$ | 5pp variation<br>Only 2pp if $T_{2,min} > TE$, but underestimated | Shorter $\Delta TE$ can improve consistency in MWF estimation |
| **Summary** | $nT_2 = 40$ – reasonable unless significant signal variation or more peaks are expected at short $T_2$<br>$\chi^2/\chi_{min}^2 = 1.02$ – SNR dependent, should typically not be chosen higher<br>gm $T_2$ times – most accurate, if $T_{2,min} \leq MW T_2$, but with MW $T_2$ unknown<br>MWF – most accurate, if $T_{2,min} \leq TE_1$, shorter within a few ms (1 – 2 ms) |  |  |  |  |

## Conclusions

The selection of shorter echo spacing and an appropriate T_2,*min*_ improved the estimation of the amount of short T_2_ signal. We showed by realistic simulations that voxel-based averages were able to estimate the true MWF within 0.13 – 4 percentage points, with increasing accuracy if T_2,*min*_ was selected slightly shorter than the first echo time. The number of T_2_ times used to compute the T_2_ distributions hardly affected the results. By contrast, increasing regularization should be applied with caution. Reporting the estimated SNR and average FA for specific regions of interest may help researchers to assess the systematic underestimation in MWF incurred due to the dependence of the T_2_ signal decay on noise and B1^+^ inhomogeneities. Advances in MR hardware may overcome current limitations in multi-echo T_2_ acquisition schemes to map the MWF more accurately.

## ACKNOWLEDGMENTS

This work was funded by the Natural Sciences and Engineering Research Council of Canada and the National Multiple Sclerosis Society. AR is funded by Canada Research Chairs.

---

**Fig. 8.**
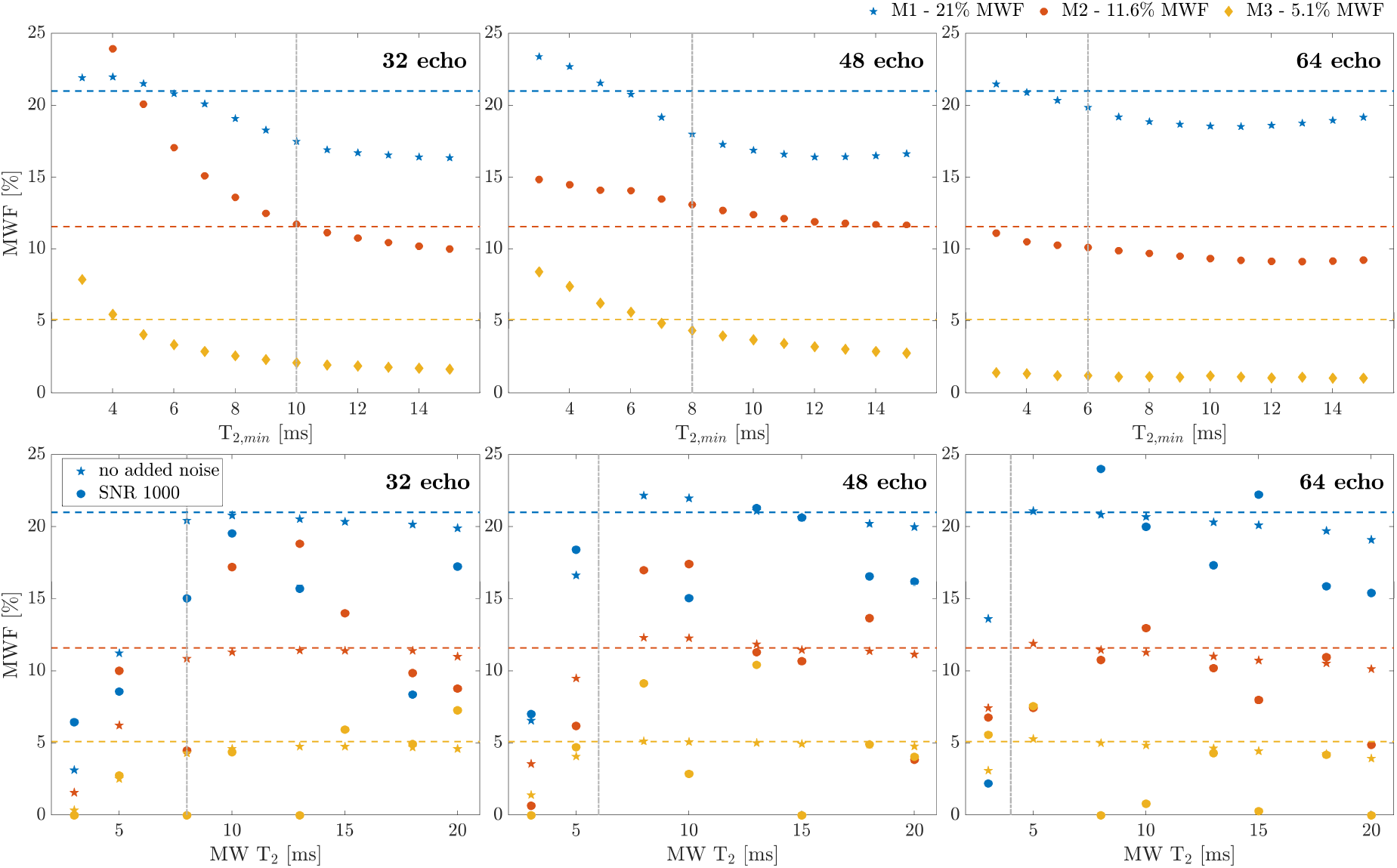
Supplementary Figure (full version of Figure 7): Mean MWF estimates relative to the true values at different T_2,min_ (top) and MW T_2_ times (bottom), at SNR = 1000, 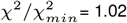, nT2= 120. Top row: Mean MWF averaged over different MW T_2_ times (15, 13, 10 and 8 ms) for different acquisition strategies at FA = 170*°*. The true MWF values (21, 11.6 and 5.1%) are shown for comparison as dashed lines. The gray, dash-dotted verticals indicate the acquisition TE, relative to T_*2,min*_. The 32e acquisition showed greater variations with T_*2,min*_ than the other acquisitions. Bottom row: Mean MWF values for different MW T_2_ times computed at a specified T_*2,min*_ (gray, dash-dotted vertical line). Estimates are compared in the presence and absence of noise. Note that the x-axes differ; top row T2,min, bottom row MW T_2_.

**Table 4.** Supplementary table: Influence of different FAs and T_*2,min*_ on the estimation of IEW gmT_2_ for all models using a MW T_2_ of 15 ms (upper section) and MW T_2_ of 8 ms (bottom section). Values represent averages are over all nT_2_ and *χ*^2^ = 1.02. Values shown are for Model 1 (21% MWF) / Model 2 (11.6%) / Model 3 (5.1%) and the different acquisition strategies.

| $T_{2,min}$ [ms] | MW gm$T_2$ [ms]<br>(true value: $T_2 = 20$ ms) | | | | | MWF [%]<br>(true values: 21%, 11.6%, 5.1%) | | | | |
| --- | --- | --- | --- | --- | --- | --- | --- | --- | --- | --- |
|  | 5 | 8 | 10 | 13 | 15 | 5 | 8 | 10 | 13 | 15 |
| M1 - 32e | 66.85 | 65.88 | 66.59 | 67.86 | 68.74 | 68.36 | 68.45 | 68.21 | 68.47 | 69.27 |
| M1 - 48e | 65.80 | 65.85 | 66.21 | 67.44 | 68.00 | 66.82 | 66.59 | 66.84 | 67.12 | 67.70 |
| M1 - 64e | 68.68 | 68.59 | 68.59 | 68.73 | 68.64 | 67.42 | 66.92 | 67.13 | 67.35 | 67.68 |
| M2 - 32e | 68.57 | 68.76 | 69.23 | 69.76 | 70.53 | 66.52 | 66.72 | 67.40 | 68.21 | 68.90 |
| M2 - 48e | 67.24 | 67.22 | 67.54 | 68.19 | 68.45 | 64.97 | 65.13 | 65.30 | 65.71 | 65.84 |
| M2 - 64e | 65.40 | 65.60 | 65.87 | 66.12 | 66.37 | 64.29 | 64.35 | 64.30 | 64.49 | 64.70 |
| M3 - 32e | 68.78 | 68.81 | 68.95 | 69.14 | 69.32 | 68.55 | 68.63 | 68.74 | 68.49 | 68.53 |
| M3 - 48e | 65.75 | 65.70 | 65.70 | 65.70 | 65.70 | 68.60 | 68.65 | 68.66 | 68.67 | 68.66 |
| M3 - 64e | 66.41 | 66.53 | 66.81 | 66.37 | 66.47 | 68.66 | 68.57 | 68.46 | 68.51 | 68.49 |
| $T_{2,min}$ [ms] | MW gm$T_2$ [ms]<br>(true value: $T_2 = 13$ ms) | | | | | MWF [%]<br>(true values: 21%, 11.6%, 5.1%) | | | | |
|  | 5 | 8 | 10 | 13 | 15 | 5 | 8 | 10 | 13 | 15 |
| M1 - 32e | 65.11 | 65.77 | 66.33 | 67.10 | 67.51 | 70.98 | 71.27 | 71.50 | 72.01 | 72.02 |
| M1 - 48e | 69.56 | 70.15 | 70.52 | 71.26 | 71.49 | 68.63 | 68.84 | 69.34 | 70.22 | 70.82 |
| M1 - 64e | 68.74 | 69.85 | 70.44 | 71.24 | 71.67 | 68.43 | 69.44 | 70.12 | 70.94 | 71.49 |
| M2 - 32e | 67.31 | 67.34 | 67.37 | 67.75 | 67.61 | 69.21 | 69.44 | 69.59 | 69.99 | 69.99 |
| M2 - 48e | 68.24 | 68.92 | 69.30 | 70.03 | 70.03 | 66.72 | 66.78 | 66.77 | 66.96 | 66.81 |
| M2 - 64e | 68.64 | 68.68 | 69.07 | 69.73 | 69.72 | 67.19 | 67.47 | 67.73 | 68.15 | 68.23 |
| M3 - 32e | 68.21 | 68.17 | 68.15 | 68.18 | 68.14 | 70.74 | 70.88 | 71.24 | 71.26 | 71.74 |
| M3 - 48e | 71.12 | 71.27 | 71.41 | 71.36 | 71.25 | 69.72 | 69.76 | 69.78 | 69.75 | 69.81 |
| M3 - 64e | 67.95 | 67.94 | 67.91 | 67.93 | 67.91 | 69.73 | 69.79 | 69.93 | 70.03 | 69.88 |

